# Dynamic interactions of type I cohesin modules fine-tune the structure of the cellulosome of Clostridium thermocellum

**DOI:** 10.1101/328088

**Authors:** Anders Barth, Jelle Hendrix, Daniel Fried, Yoav Barak, Edward Bayer, Don C. Lamb

## Abstract

Efficient degradation of plant cell walls by selected anaerobic bacteria is performed by large extracellular multienzyme complexes termed cellulosomes. The spatial arrangement within the cellulosome is organized by a protein called scaffoldin, which recruits the cellulolytic subunits through interactions between cohesin modules on the scaffoldin and dockerin modules on the enzymes. Although many structural studies of the individual components of cellulosomal scaffoldins have been performed, the role of interactions between individual cohesin modules and the flexible linker regions between them are still not entirely understood. Here, we report single-molecule measurements using Förster resonance energy transfer to study the conformational dynamics of a bimodular tandem cohesin segment of the scaffoldin protein CipA of *Clostridium thermocellum*. Our data reveal the existence of compacted structures in solution that persist on the timescale of milliseconds. The compacted conformation is found to be in dynamic equilibrium with an extended state that shows distance fluctuations on the microsecond timescale. Shortening of the inter-cohesin linker does not significantly alter the structural dynamics. Upon addition of dockerin-containing enzymes, an extension of the flexible state is observed but the cohesin-cohesin interactions persist. This suggests that the dockerin-binding interfaces are not involved in cohesin-cohesin interactions. The formation of cohesin-cohesin interactions is also observed in all-atom molecular dynamics simulations of the system. From the simulations, we identify possible inter-cohesin binding modes, none of which show obstruction of the cohesin-dockerin binding interfaces. Our results go beyond the view of scaffoldin as “beads on a string”. We propose that both the flexibility and cohesin-cohesin interactions are important factors for the precise spatial arrangement of the enzymatic subunits in the cellulosome that leads to the high catalytic synergy in these assemblies. Hence, the flexibility of the linker region and cohesin-cohesin interactions should be considered when designing cellulosomes for industrial applications.

## Introduction

Cellulose from plant cell walls is the most abundant source of renewable carbon (1). As the world reserve of fossil fuels is being depleted the conversion of plant biomass into bioethanol is a promising approach to solve the global energy problem. The efficient degradation of plant cell wall material, however, remains a challenge due to the hydrolytic stability of cellulosic polysaccharides (2–4). In nature, aerobic bacteria and fungi secrete specialized enzymes to break down cellulose into oligosaccharides. In contrast, anaerobic bacteria utilize a cell-attached extracellular megadalton multienzyme complex, called the cellulosome, to efficiently degrade plant cell walls (5–7). The spatial proximity of cellulases and hydrolases within the cellulosome results in its highly synergetic catalytic activity (8).

The cellulosome of the thermophilic anaerobe *Clostridium thermocellum* contains the extracellular scaffoldin protein CipA that mediates the cell-substrate interactions via the cellulose binding module (CBM) (Fig. 1A) (9–12). Cell wall attachment is achieved through the binding of a type II dockerin (DocII) module in CipA to a type II cohesin (CohII) module in a cell-surface associated protein linked to the cell through a surface-layer homology domain (SLH). An X module adjacent to the DocII module has been shown to have an important function for the binding interaction (13). CipA contains a linear array of nine type I cohesin (CohI) modules with the CBM located between CohI 2 and 3. Each CohI module acts as an attachment site for various cellulose-processing enzymes which bind with high affinity through their type I dockerin (DocI) modules (14–16). The interactions between the CipA CohI and the enzyme-borne DocI modules of different enzymes are non-specific and thus allow for variability in the composition of the catalytic subunits (17, 18).

**Figure 1:**
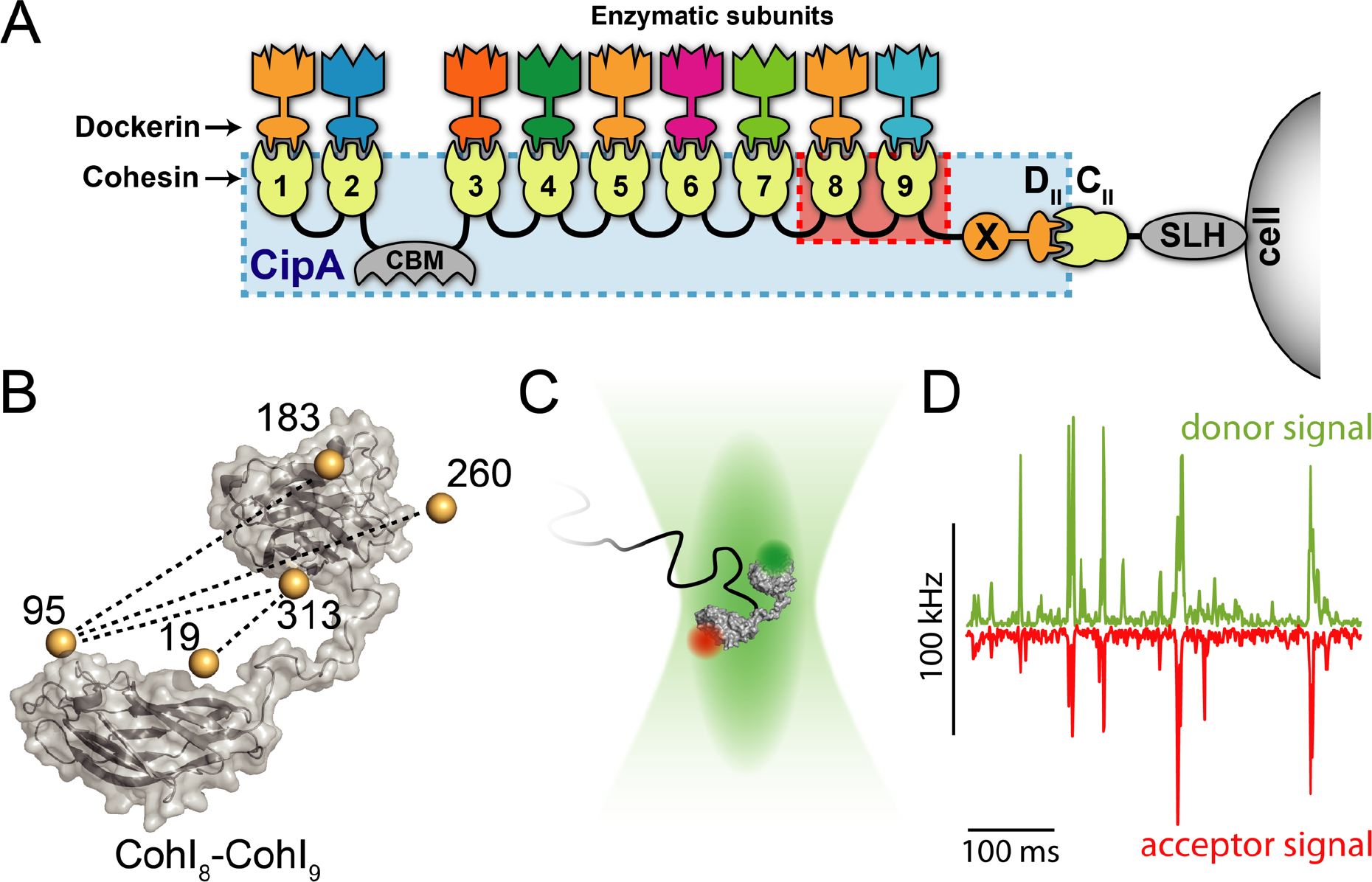
Structure of the cellulosome of *C. thermocellum*. A) The cellulosome complex is composed of the scaffoldin protein (CipA, blue box) that is anchored to the cell surface and the cellulose, offering binding sites for cellulose-processing enzymes. Surface attachment occurs through a surface-layer homology domain (SLH) linked to a type II cohesin module (C_II_), that interacts with a type II dockerin module (D_II_) connected to an X module (X) at the C-terminus of CipA. Binding to cellulose occurs *via* the cellulose binding module (CBM). CipA contains nine type I cohesin modules (1–9), that can each bind to a type I dockerin module on different cellulose-processing enzymes. In this work, the CohI_8_-CohI_9_ fragment of CipA is studied (highlighted in red). B) Atomistic model of the CohI_8_-CohI_9_ fragment defined from a homology model. Bronze spheres indicate the average fluorophore positions determined from geometrical calculations of the accessible volume(36). Dashed lines show the labeling combinations investigated by single-molecule FRET. C) In the experiment, fluorescently-labeled molecules are measured as they freely diffuse through the confocal volume. D) Singlemolecule events result in coinciding bursts of fluorescence in the donor and acceptor detection channels.

Previous x-ray crystallography and NMR spectroscopy studies revealed the structure of many individual components of the cellulosome (19, 20). However, due to the inherently dynamic quaternary structure of the cellulosome (21, 22), the precise arrangement of the components linked by the scaffoldin protein remains poorly understood. Low-resolution structural methods such as small angle x-ray scattering (SAXS) have revealed a dynamic picture of different artificial (23) and natural scaffoldin (24–26) fragments. Additional structural insights were obtained by cryo-electron microscopy (cryoEM) on a fragment of CipA consisting of three consecutive CohI modules, which in contrast showed a compacted structure with outward pointing catalytic domains (27). The *C. thermocellum* scaffoldin segments are connected by flexible linkers that are 20 to 40 residues long and rich in proline and threonine residues. These inter-cohesin linkers were found to be predominantly disordered by molecular dynamics (28) and SAXS (24–26) studies, but were proposed to adopt a predominantly extended structure based on recent NMR data (29). The structural flexibility may be essential for the efficient access to the crystalline cellulose substrate within the heterogeneous environment of the plant cell wall containing hemicellulose, lignin and pectin components. The connection of cohesins by linkers has been reported to enhance the catalytic activity in mini-cellulosome model systems by factor of ~2, however contradictory results have been obtained regarding the effect of linker length and composition (23, 30).

To directly measure the dynamics of scaffoldin and thereby investigate the role dynamics play for the cellulosome, we investigated the conformational dynamics of a tandem fragment of CipA consisting of the cohesin I modules 8 and 9 connected by the 23-residue long wildtype linker (Fig. 1A, red square, and Fig. 1B) using single-molecule Förster resonance energy transfer (smFRET). The CohI_8_-CohI_9_ fragment underwent transitions between compacted and extended structures on the millisecond timescale. Quantitative information about the interconversion rates was obtained from dynamic photon distribution analysis (dynamic PDA) (31, 32) and filtered fluorescence correlation spectroscopy (fFCS) (33). The effect of the linker on the structural dynamics was probed by shortening of the linker peptide, and the influence of the CohI-DocI interactions was investigated using the enzymes Cel8A and Cel48S. We also complemented the experimental data with all-atom molecular dynamics (MD) simulations to gain further insights into possible inter-cohesin binding modes on the atomic level.

## Results and Discussions

### Single-molecule FRET identifies interactions in the CohI_8_-CohI_9_ fragment

We investigated the inter-modular distance fluctuations of the CohI_8_-CohI_9_ fragment from CipA using a single-molecule FRET (smFRET) analysis (34). In this method, fluorescently labeled single molecules are measured at picomolar concentrations in solution as they diffuse through the observation volume of a confocal fluorescence microscope on the timescale of ~1 ms (Figure 1 C-D). For every single molecule event, the efficiency of energy transfer from a donor fluorophore to an acceptor fluorophore reports on the inter-dye distance and thus the conformation of the molecule. In contrast to ensemble methods, which report an average value, smFRET is particularly suited to study the conformational heterogeneity of biomolecules by measuring intramolecular distances one molecule at a time. SmFRET experiments were performed for different constructs probing a total of four distances between the CohI modules. For fluorescent labeling, we introduced cysteines at positions 19, 95, 183, 260 and 313 in the CohI_8_-CohI_9_ fragment to obtain the four combinations of labeling positions 19-313, 95-313, 95-183 and 95-260 (Fig. 1B). The attachment sites of the fluorophores were chosen outside of the dockerin binding interface (Fig. S1) (35). For each CohI_8_-CohI_9_ mutant, these positions were stochastically labeled with the dyes Atto532 and Atto647N. No local influence of the labeling position on the photophysical properties of either fluorophore was observed, allowing quantitative analyses without the necessity for site-specific labeling. The results of the smFRET experiments are shown in Fig. 2 A-D. The molecule-wise FRET efficiency histograms are shown together with the fit to three-component Gaussian distributions. All constructs showed a major FRET population with low to intermediate FRET efficiency (red dashed lines) and a high FRET efficiency population (E > 0.8, blue dashed lines). An intermediate FRET efficiency population (yellow dashed lines) connects the two populations. In construct 19-313, no intermediate population was detected by the analysis, but rather a small low FRET efficiency population was observed (purple dashed line).

**Figure 2:**
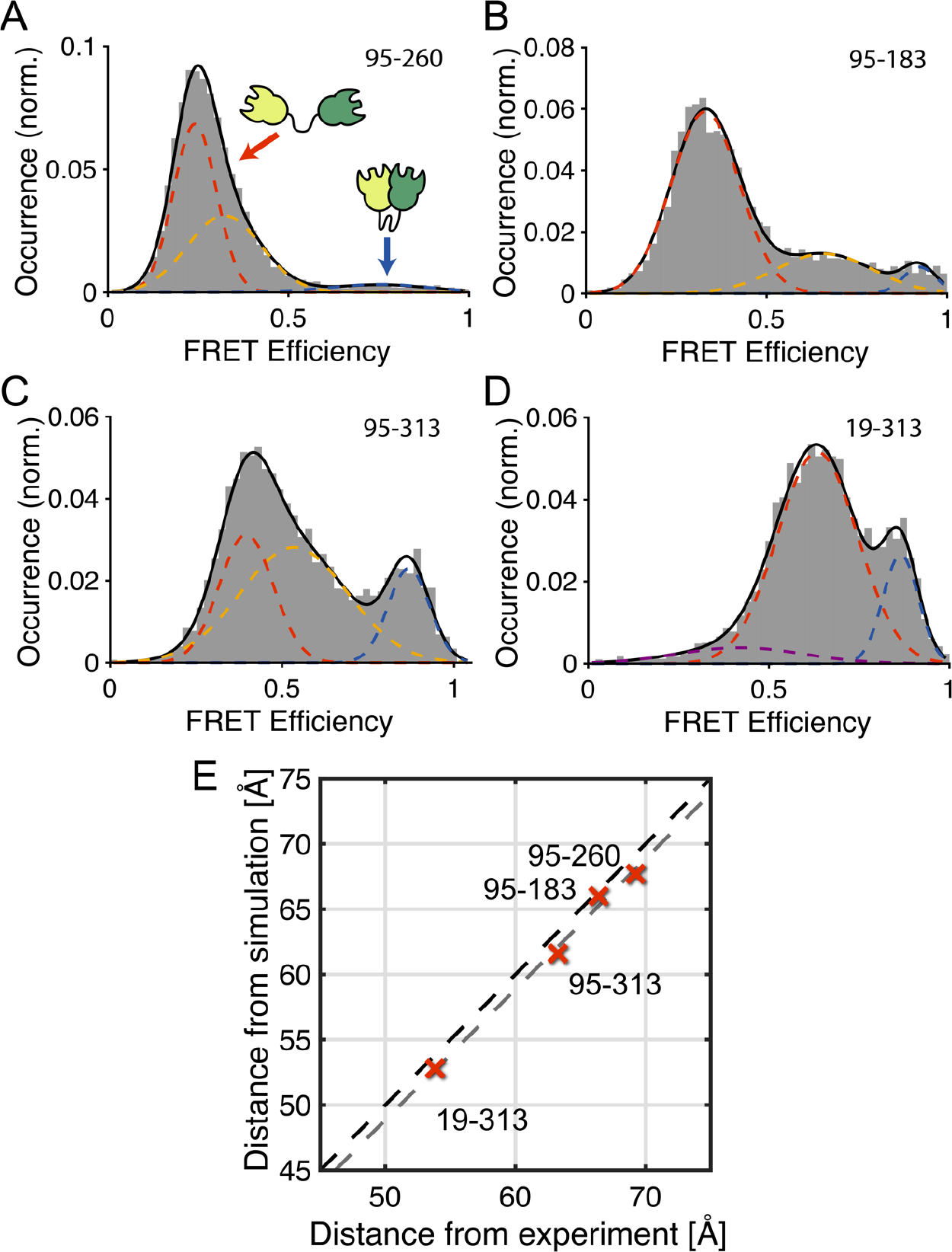
Conformations of the investigated CohI_8_-CohI_9_ fragment. A-D) Single-pair FRET efficiency histograms for different combinations of labeling positions as indicated in Figure 1B. Respective residues were mutated to cysteines and labeling was performed stochastically with the dyes Atto532 and Atto647N. Histograms were fit using three Gaussian distributions, revealing low (red), medium (yellow) and high (blue) FRET efficiency populations. In the 19-313 construct (D), no medium FRET efficiency population is observed, but an additional low FRET efficiency population is detected (purple). E) A comparison of the measured and simulated distances for the different constructs. A linear correlation is observed between the distances determined from the main population (dashed red lines in A-D) of the single-pair FRET experiments and from rigid-body torsion-angle molecular dynamics simulations. The experimental FRET efficiencies were converted into distances using a Förster radius of 59 υ. The black dashed line indicates a linear correlation, while the grey dashed line indicates linear correlation with an offset of 1.1 υ, as determined from the average deviation between experimental and theoretical distances.

To estimate expected FRET efficiencies of the dynamic, non-interacting CohI_8_-CohI_9_ fragment for all tested mutants, we performed simplified molecular dynamics (MD) simulations in the absence of solvent, treating the CohI modules as rigid bodies. Under the conditions of the simulation, indeed no stable interactions occurred between the CohI modules, as is evident from the root-mean-square deviation (RMSD) of the MD trajectory which shows vanishing correlation after ~100 ps (Fig. S2 A-B). From the simulation, average FRET efficiency values are extracted using accessible volume calculations to determine sterically accessible positions for the dyes at every time step (36). We obtained good agreement between smFRET results for the low-FRET conformation and the simulated MD data (R^2^ > 0.98, Fig. 2E and Table 1) with an average deviation of 1.1 Å. The discrepancy is within the absolute experimental error one would expected for smFRET measurements (37). The good agreement between smFRET results and the MD data suggests that the “open” conformation, characterized by the absence of long-lived interactions between the Cohl modules, corresponds to the dominant CohI_8_-CohI_9_ conformation.

**Table 1:**
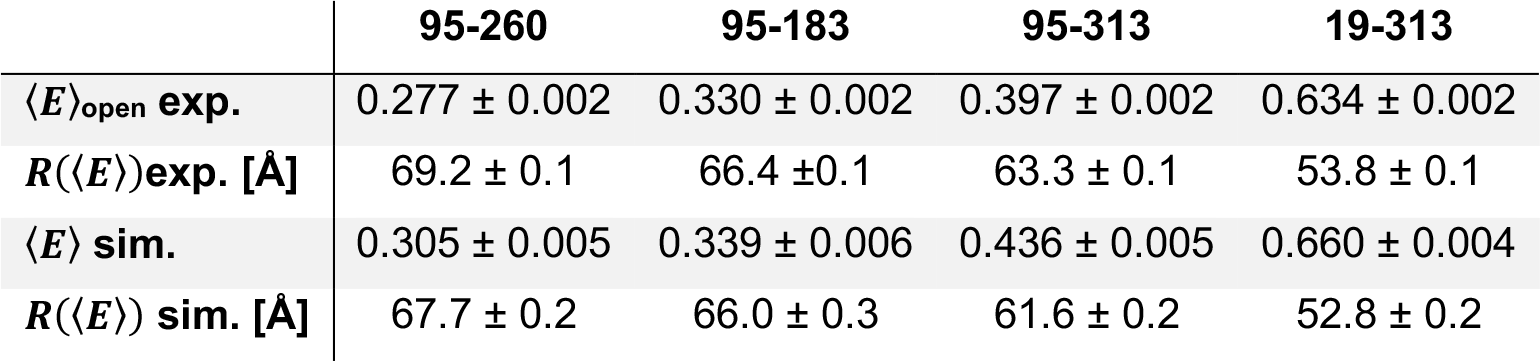
Comparison of experimental results and molecular dynamics simulations of the extended conformation. Mean FRET efficiencies 〈*E*〉 of the open state are obtained from fits of the experimental distributions of corrected FRET efficiencies with a three component Gaussian mixture model, and converted to distances (Förster radius R_0_ = 59 Å). For simulations, the FRET efficiency is calculated at every time step, averaged over the total trajectory and then converted into a distance *R*(〈*E*〉) (see M&M).

|  | 95-260 | 95-183 | 95-313 | 19-313 |
| --- | --- | --- | --- | --- |
| $\langle E \rangle_{\text{open exp.}}$ | $0.277 \pm 0.002$ | $0.330 \pm 0.002$ | $0.397 \pm 0.002$ | $0.634 \pm 0.002$ |
| $R(\langle E \rangle)_{\text{exp. [Å]}}$ | $69.2 \pm 0.1$ | $66.4 \pm 0.1$ | $63.3 \pm 0.1$ | $53.8 \pm 0.1$ |
| $\langle E \rangle_{\text{sim.}}$ | $0.305 \pm 0.005$ | $0.339 \pm 0.006$ | $0.436 \pm 0.005$ | $0.660 \pm 0.004$ |
| $R(\langle E \rangle)_{\text{sim. [Å]}}$ | $67.7 \pm 0.2$ | $66.0 \pm 0.3$ | $61.6 \pm 0.2$ | $52.8 \pm 0.2$ |

Since previous studies revealed a highly dynamic structure for the related CohI_1_-CohI_2_ fragment (26), we expected all CohI_8_-CohI_9_ mutants to show a FRET efficiency distribution with a single peak for all constructs, as would be expected from fast dynamic averaging over many conformations on the timescale of diffusion through the confocal volume of ~1 ms. The existence of a high FRET efficiency population in all four constructs suggests specific interactions between the two Cohl modules that persist on the millisecond timescale. The observation of a characteristic bridge between the high FRET efficiency (“closed”) population and the low-to-intermediate FRET efficiency (“open”) conformation is additionally indicative of dynamic interconversion between the states. For the 19-313 construct, no intermediate FRET efficiency population is detected, likely because the difference between the FRET efficiencies of the open (E ≈ 0.6) and closed (E ≈ 0.9) is not large enough to resolve the conformational dynamics. The rigid-body MD simulations showed no stable interactions between the Cohl modules and thus sampled the dynamic state of CohI_8_-CohI_9_, agreeing well with the experimentally determined distances of the main population.

### The CohI_8_-CohI_9_ fragment shows conformational dynamics on the millisecond timescale

To further investigate the conformational dynamics of CohI_8_-CohI_9_, we employed the FRET-2CDE filter that identifies FRET fluctuations based on anti-correlated changes of the donor and FRET-sensitized acceptor signals (Fig. 3A) (38). The FRET-2CDE filter is defined such that a value of 10 signifies no dynamics, while any larger values indicate the presence of conformational transitions on the time scale between 0.1 to 10 milliseconds. The two-dimensional plot of FRET efficiency against FRET-2CDE filter (Fig. 3A) revealed a systematic deviation from the static line at a value of 10 for the FRET-2CDE filter for single-molecule events with intermediate FRET efficiency. We also examined the fluorescence lifetime of the donor fluorophore to further investigate the dynamics of the CohI_8_-CohI_9_ construct (Fig. 3B) (32). The lifetime of the donor fluorophore offers an independent read-out of the FRET efficiency. In a plot of the FRET efficiency against the donor fluorescence lifetime (Fig. 3B), one can define a static FRET line (green line in Fig. 3B) that single-molecule events showing no conformational dynamics will lie on. A systematic shift towards longer fluorescence lifetimes results when conformational transitions occur during a single-molecule event. In the case of a two-state dynamic system, the dynamic FRET line can be calculated by considering all mixtures of the two states (dashed red line in Fig. 3B). The data clearly showed a systematic deviation from the static FRET line that could be explained by two-state conformational dynamics with a lifetime value for the donor fluorophore of 0.6 ns and 2.9 ns. The same qualitative result was obtained for all four constructs (Fig. S3 and Table S1). Interestingly, the low-FRET state was also shifted from the dynamic FRET line, although it showed a value around 10 for the FRET-2CDE filter. Since the FRET-2CDE filter relies on a kernel density estimator, effectively smoothing over a finite time window (here 100 μs), it is not sensitive to faster fluctuations. The lifetime-based readout, however, is independent of the timescale of the dynamics since it only relies on the mixing of different conformations during a singlemolecule event. This suggests that the extended state is highly dynamic on the microsecond timescale, faster than the averaging window of 100 μs chosen for the calculation of the FRET-2CDE filter. As a control, we measured a construct where both the donor and acceptor dyes were placed on the CohI_9_ module (Fig. S4). As expected, the control construct showed a single population and exhibits no deviation from the static FRET line.

**Figure 3:**
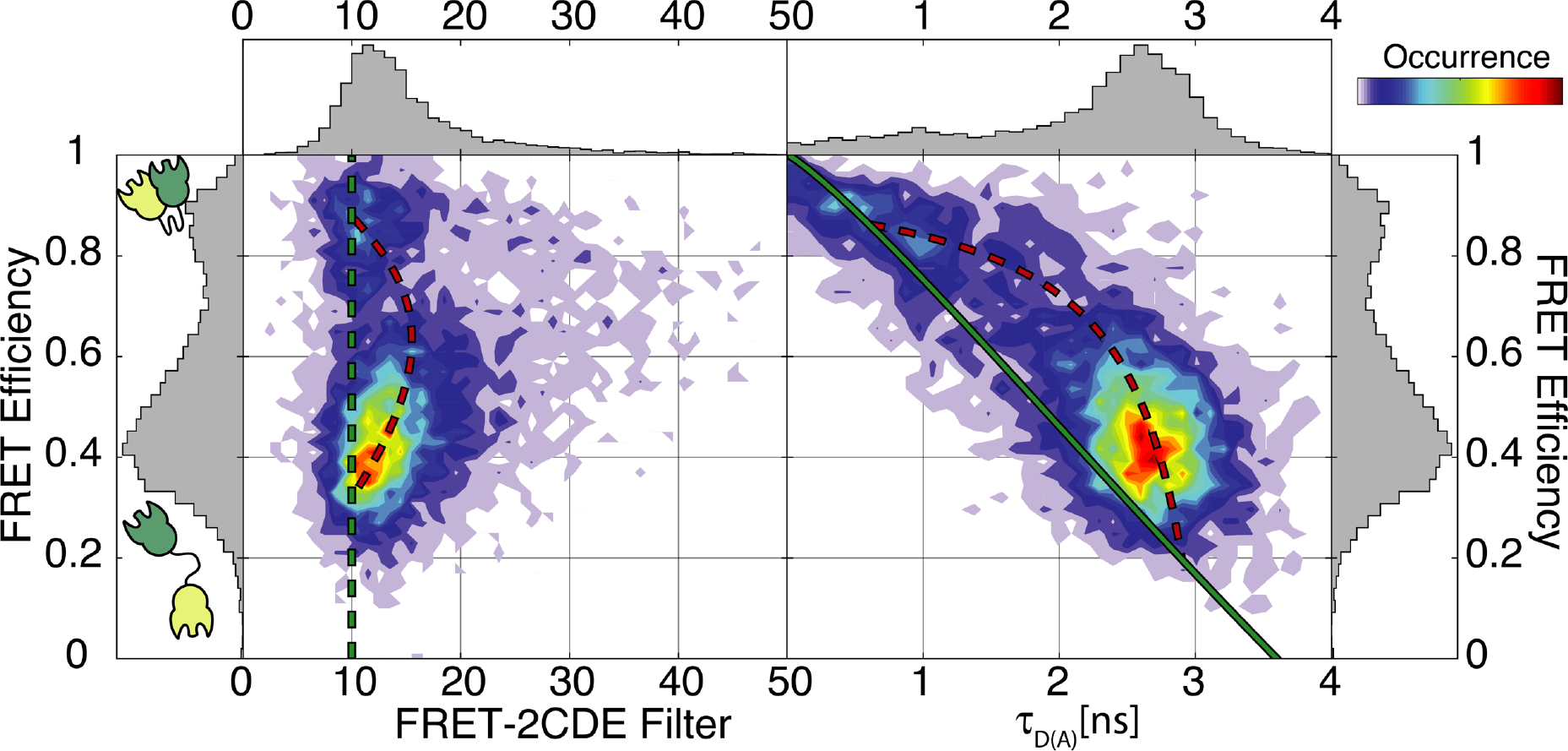
Identification of conformational dynamics in the CohI_8_-CohI_9_ construct 95-313. Left: 2D histogram of the FRET-2CDE filter versus FRET efficiency. Values larger than 10, visible for events with intermediate to high FRET efficiency, indicate dynamics within the burst. Right: 2D histogram of the donor lifetime versus FRET efficiency. A systematic shift from the static FRET line (green) is observed, consistent with the presence of conformational dynamics. Additionally, the low-to-intermediate FRET efficiency species at ~40 % also deviates from the static FRET line, indicating fast dynamics on the microsecond timescale. The dynamic behavior can be described theoretically using the dynamic FRET line (dashed red line). The start and end points were extracted from fluorescence decay analysis of all molecules using a biexponential model function (Table S1).

The qualitative assessment of the conformational dynamics in CohI_8_-CohI_9_ confirms the hypothesis of a compacted state which is in dynamic equilibrium with an open, flexible state. Additional information is obtained about the timescales of the dynamic processes. Transition to and from the compacted state occur on the millisecond timescale, while the open state exhibits fast fluctuations on the microsecond timescale. In the following sections, we focus on the characterization and quantification of the dynamics.

### Photon distribution analysis quantifies the dynamics between open and closed states

To quantify the timescale of the dynamics, we utilized additional analysis methods. First, we focus on the millisecond dynamics between the open and closed states. Photon distribution analysis (PDA) is a powerful tool to disentangle the contributions of photon shot noise to the width of the observed FRET efficiency distribution, from physically relevant factors such as static conformational heterogeneity (31). In its simplest form, PDA assumes a Gaussian distribution of distances that is transformed using the known photon statistics of the measurement and the experimental correction factors to obtain the corresponding shot-noise limited proximity ratio (*PR*) histogram. PDA can also be used to describe the effect of conformational dynamics on the observed proximity ratio histogram (32). By sectioning the photon counts into equal time intervals, the resulting proximity ratio histogram can be described analytically using a two-state kinetic model, whereby each individual state’s heterogeneity is described by a distribution of distances. To increase the robustness of the analysis, each dataset was processed using different time window sizes (0.25 ms, 0.5 ms and 1 ms) and globally fit with respect to the interdye distances and kinetic rates.

We performed dynamic PDA of all different constructs (Fig. 4 and Table 2). In addition to the two interconverting species, a static low-FRET population was needed to account for the contributions of small amounts (1-5 %) of low FRET efficiency species likely caused by acceptor blinking. The kinetic model was successful in describing the observed distributions of FRET efficiencies in all cases (Fig. S5). While for the 19-313 construct no intermediate-FRET population was detected from the analysis of the FRET efficiency distribution (Fig. 2D), the global dynamic PDA was successful in recovering kinetic information for this construct (Fig. 4D). In all construct, the extended state was more populated than the closed state, suggesting a fast rate of opening and a slower rate of contact formation. This is confirmed by the extracted rates for closing of *k*_oc_ = 0.5 ± 0.3 ms^−1^ and opening of *k*_co_ = 2.0 ± 0.4 ms^−1^, corresponding to dwell time of ~2 ms in the open state and ~0.5 ms in the closed state. The determined center distances from PDA are in good agreement with the previous analysis of the FRET efficiency histograms, deviating by less than 3 Å (Table 1 and 2).

**Figure 4:**
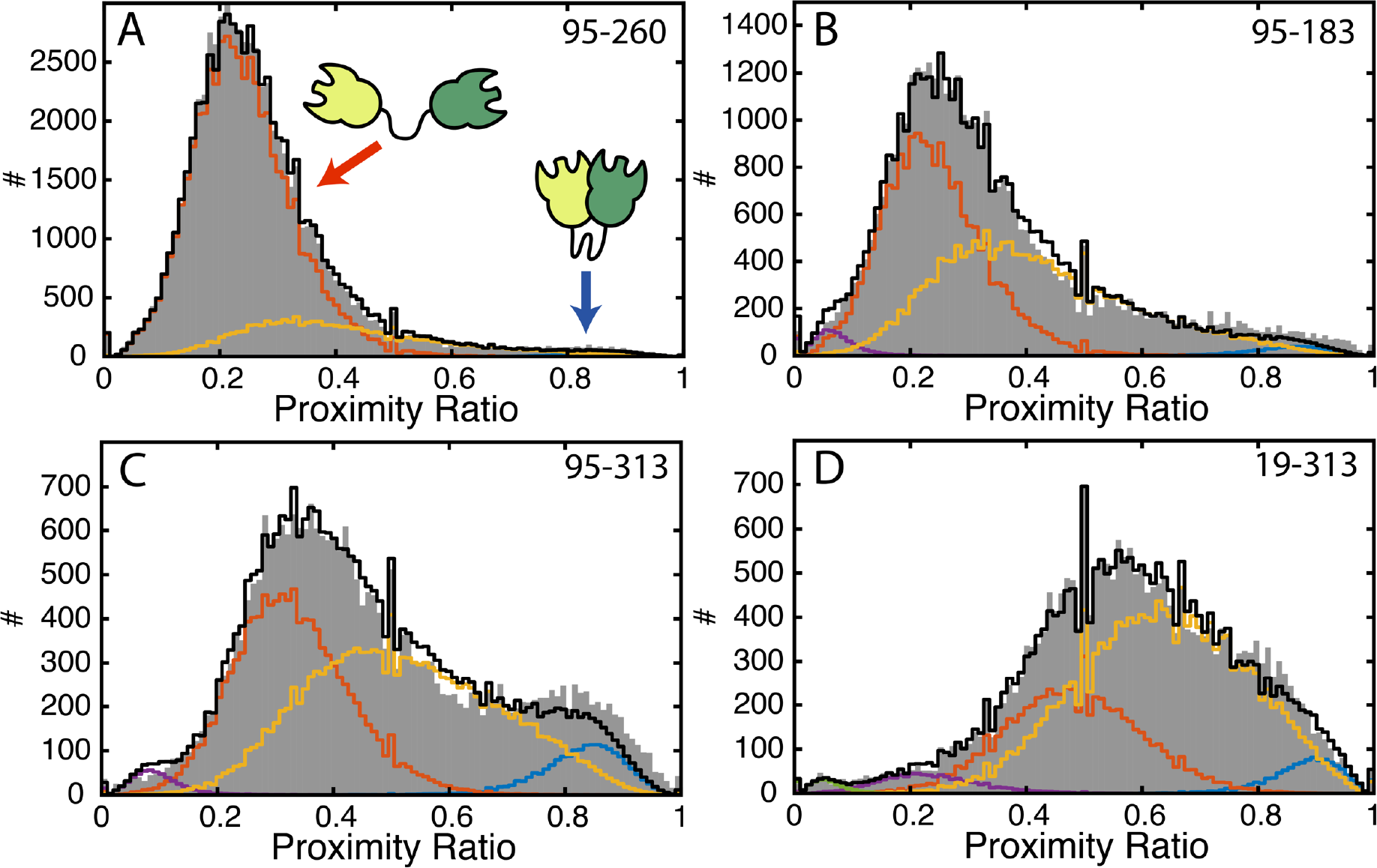
Dynamic photon distribution analysis (PDA) of the wildtype CohI_8_-CohI_9_ fragment. To investigate the timescale of the indicated dynamics, proximity ratio histograms for constructs 95-260 (A), 95-183 (B), 95-313 (C) and 19-313 (D) were fit using dynamic PDA. Shown are the data for a time window size of 1 millisecond. The proximity ratios (uncorrected FRET efficiency) are shown as grey bars and the dynamic PDA fits are shown as black lines. In addition, the low (red) and high (blue) FRET efficiency populations and the population of molecules showing interconversion during the observation time (yellow) are indicated. Additional static low-FRET efficiency populations are shown in purple and green. See Fig. S5 for the global fit of the data using time windows of Δ*t* = 0.25 ms, 0.5 ms and 1 ms.

**Table 2:** Dynamic photon distribution analysis of the different CohI_8_-CohI_9_ constructs with wildtype (wt) linker and shortened linker. Construct 95-260 with the shortened linker showed artifacts in the measurement and was thus excluded from the analysis (see Fig. S10). Errors are given as 95% confidence intervals as determined from the curvature of the 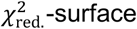.

|  | 95-260 | 95-183 | 95-313 | 19-313 |
| --- | --- | --- | --- | --- |
|  | wt linker |  |  |  |
| $R_{\text{closed [Å]}}$ | $40.2 \pm 0.6$ | $40.7 \pm 1.0$ | $41.3 \pm 0.8$ | $39.7 \pm 1.0$ |
| $R_{\text{open [Å]}}$ | $69.4 \pm 0.1$ | $68.6 \pm 0.3$ | $62.8 \pm 0.3$ | $55.6 \pm 0.6$ |
| $k_{\text{close} \rightarrow \text{open [ms}^{-1}\text{]}}$ | $2.09 \pm 0.10$ | $2.39 \pm 0.13$ | $1.41 \pm 0.09$ | $1.92 \pm 0.17$ |
| $k_{\text{open} \rightarrow \text{closed [ms}^{-1}\text{]}}$ | $0.14 \pm 0.01$ | $0.51 \pm 0.03$ | $0.57 \pm 0.05$ | $0.87 \pm 0.13$ |
|  | shortened linker |  |  |  |
| $R_{\text{closed [Å]}}$ | - | $40.0 \pm 0.6$ | $39.1 \pm 1.6$ | $38.5 \pm 0.7$ |
| $R_{\text{open [Å]}}$ | - | $66.0 \pm 0.2$ | $62.6 \pm 0.3$ | $54.7 \pm 0.3$ |
| $k_{\text{close} \rightarrow \text{open [ms}^{-1}\text{]}}$ | - | $1.94 \pm 0.10$ | $2.14 \pm 0.14$ | $1.79 \pm 0.23$ |
| $k_{\text{open} \rightarrow \text{closed [ms}^{-1}\text{]}}$ | - | $0.37 \pm 0.02$ | $0.67 \pm 0.05$ | $0.38 \pm 0.06$ |

### Correlation analysis identifies a second kinetic state on the microsecond timescale

Secondly, we looked for conformational dynamics on the microsecond timescale. To this end, we applied fluorescence correlation spectroscopy (39, 40). As a first step, we calculated the auto- and cross-correlation functions of the donor fluorescence and the FRET-induced acceptor signal. In the presence of conformational dynamics, the FRET fluctuations cause an anti-correlation contribution to the cross-correlation function and matching positive contributions to the autocorrelation functions. Representative auto- and cross-correlation functions for construct 95-260 are shown in Fig. 5A and for all constructs in Fig. S6. From the FRET-FCS analysis, we obtained a kinetic relaxation time of 24 ± 7 μs across all four constructs (Table 3). To obtain a more detailed picture of these fast dynamics, we applied filtered-FCS (fFCS) (33, 41). Filtered-FCS uses the lifetime, anisotropy and color information available for each photon to assign statistical weights (or filters) with respect to two or more defined species (Fig. S7). To define the open and closed conformation, the characteristic patterns were determined from the measurement directly by pooling data from the low or high FRET efficiency events (Fig S7B). The inclusion of additional dimensions and the application of statistical weighting in fFCS leads to increased contrast in comparison to the FRET-FCS analysis. The fFCS crosscorrelation functions for the four constructs are shown in Fig. 5B. Fitting with a single kinetic term revealed systematic deviations in the residuals, prompting us to include a second kinetic term (Fig. S8). The extracted timescales of the two terms are 15 ± 4 μs and 600 ± 200 μs for the fast and slow component, respectively (Table 3). The timescale of the slow component overlapped with the timescale of diffusion (~1-2 ms), resulting in the relatively high uncertainty. The amplitude of the fast term with significantly higher, contributing to 79 ± 7 % of the observed dynamics. Because the relaxation time of the slow component agrees well with that determined from the dynamic-PDA analysis (τ_*R*_ = 410 ± 80 μs), we assign the slow component to the previously described slow transition between the open and closed state. The fast component identified by fFCS consequently corresponds to the dynamics of the freely fluctuating open state that occur on the timescale of ~15 μs.

**Figure 5:**
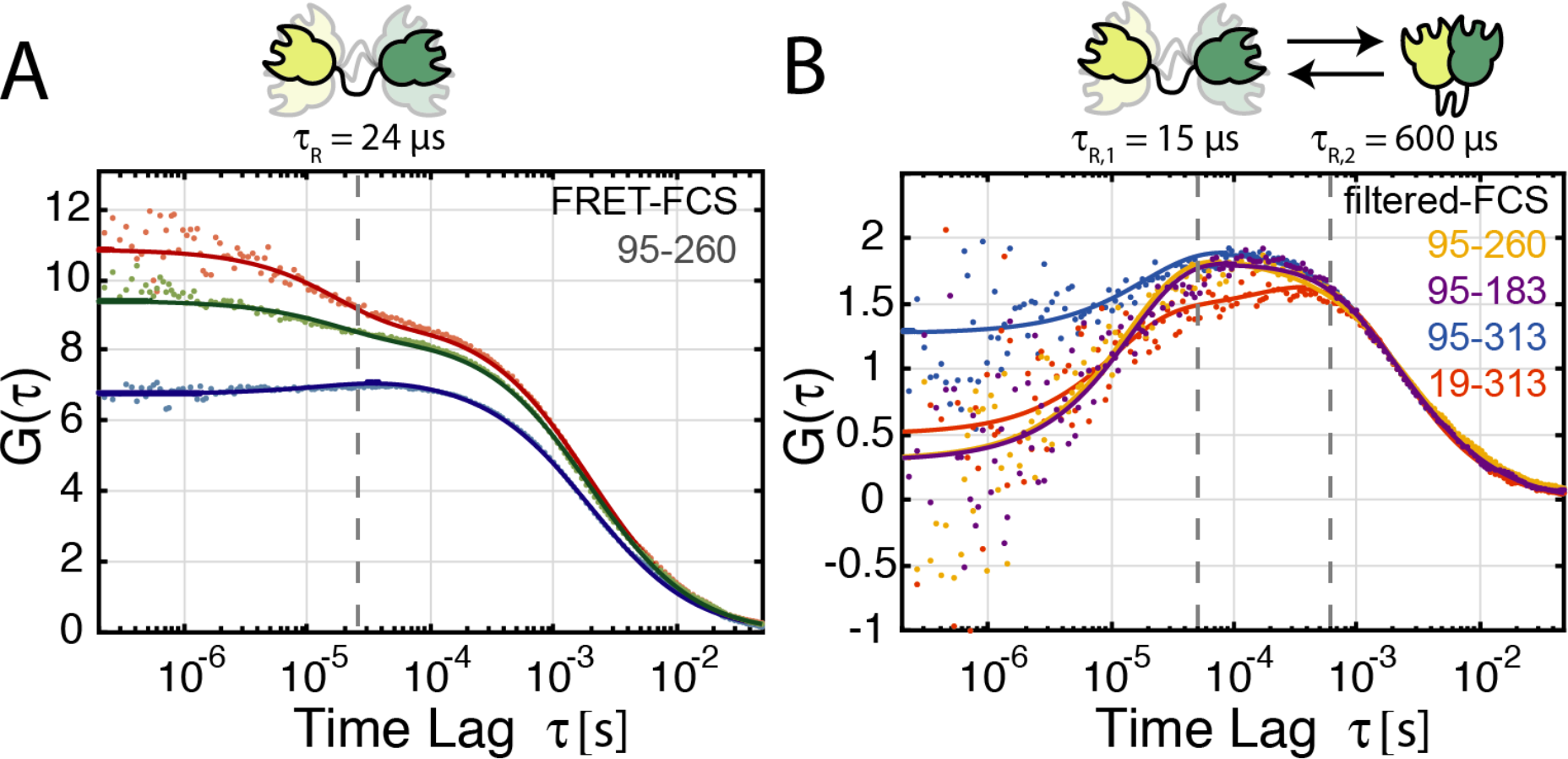
FCS analysis of conformational dynamics. A) FRET-FCS reveals a bunching term in the donor fluorescence and FRET-induced acceptor fluorescence autocorrelation curves (green and red curves), as well as a corresponding anti-correlation term in the cross-correlation function (blue curve). A global analysis reveals dynamics with a timescale of 24 ± 7 μis. Data are shown for the 95-260 construct. B) The filtered-FCS cross-correlation functions are shown for the constructs 95-260 (yellow), 95-183 (purple), 95-313 (blue) and 19-313 (red). Filtered-FCS enhances the contrast of the kinetic contributions to the correlation function. The species cross-correlation curves for the four constructs show similar timescale of dynamics, revealing two terms, one at 15 ± 4 μs and a second at 600 ± 200 μis with contributions of 76 ± 4 % and 24 ± 4 %, respectively.

**Table 3:** Species-selective fluorescence correlation spectroscopy analysis of the different CohI_8_-CohI_9_ constructs with wildtype (wt) linker and shortened linker. Given are the average and standard deviation of the relaxation times, *τ_R_*, and relative amplitudes, *f*, of independent analyses of the four constructs. All three correlation functions in FRET-FCS or four correlation functions in filtered-FCS were globally fit as described in the main text. Relative amplitudes were determined based on amplitudes of the kinetic terms in the species cross-correlation functions. For the shortened linker, the construct 95-260 was excluded due to dye artifacts (see Fig. S10). For comparison, the relaxation times from dynamic PDA calculated by *τ_R_* = (*k_c→o_ + k_o→c_*)^−1^ analysis are given, averaged over all constructs for the wt linker and constructs 95-183, 95-313 and 19-313 for the shortened linker. For the results of the FCS analysis of the different constructs, see Table S2.

|  | **FRET-FCS** | **filtered FCS** |  |  |  | **dynamic PDA** |
| --- | --- | --- | --- | --- | --- | --- |
| | $\tau_R$ [μs] | $\tau_{R,1}$ [μs] | $f_1$ [%] | $\tau_{R,2}$ [μs] | $f_2$ [%] | $\tau_R$ [μs] |
| **wt linker** | 24 ± 7 | 15 ± 4 | 79 ± 7 | 600 ± 200 | 21 ± 7 | 410 ± 80 |
| **shortened linker** | 30 ± 14 | 9 ± 7 | 76 ± 5 | 300 ± 200 | 24 ± 5 | 420 ± 60 |

Our observation of a highly dynamic structure in the open state is in agreement with previous studies on artificial chimeric minicellulosomes (23) or the CipA fragments CohI_1_-CohI_2_ and CohI_2_-CBM-CohI_3_ in complex with the enzyme Cel8A (26). On the other hand, a cryo-EM study on the fragment CohI_3_-CohI_4_-CohI_5_ in complex with Cel8A revealed predominantly compacted structures showing direct interactions between the cohesin modules with outward pointing enzymes (27), similar to the compacted state observed here. The report of a small fraction of complexes in extended conformations in that study additionally supports our observation of a dynamic equilibrium between extended and compacted structures. Cohesin-cohesin interactions were also observed in the crystal structure of the C-terminal fragment of CipA, CohI_9_-X-DocII, in complex with a CohII module, showing homodimerization mediated by intermolecular contacts between the CohI_9_ modules (42), and in the crystal structure of a CohII dyad from *A. cellulolyticus* (43).

### Shortening of the linker has a minor effect on the observed dynamics

After having characterized the structure and dynamics of the wt-linker CohI_8_-CohI_9_ fragment, we turned to investigate the role of the linker by designing a construct with a shortened linker. The wt-linker is 23 residues long and is mainly composed of polar and aliphatic amino acids with a high content of threonine (39%) and proline (22%) residues (Fig. S1A). We deleted eleven residues from the center of the linker, shortening it to 12 residues without significantly altering the peptide properties (Fig. S1B). The dynamic behavior detected for the wt-linker persists in the shortened linker constructs (Fig. S9). However, construct 95-260 in combination with the shortened linker showed dye-induced artifacts in the PIE-MFD analysis and was thus excluded from the discussion (see Fig. S10). We again performed dynamic PDA to quantify the dynamics and inter-dye distance distributions (see Table 2, Fig. 6A for construct 95-313, and Fig. S5 and S11 for all constructs). No significant distance change was detected for the closed conformation (Table 2). Surprisingly, also no significant shift to shorter distances was evident for the open conformation, although construct 95-183 showed a minor contraction from 69 Å to 66 Å (Table 2 and Figure 6B). The dynamic interconversion rates of the shortened-linker constructs exhibited no major change with respect to the wt-linker with an opening rate of 2.0 ± 0.2 ms^−1^ and a closing rate of 0.5 ± 0.2 ms^−1^. As before, we performed a filtered-FCS analysis (Table 3). In contrast to the dynamic PDA, the correlation analysis detects increased interconversion rates for the shortened linker constructs. The slow component showed a relaxation time of 300 ± 200 μs (averaged over all constructs), in good agreement with the dynamic timescale detected by dynamic PDA of 420 ± 60 μs for the shortened linker construct, but faster than what we obtained from the correlation analysis for the wt-linker (600 ± 200 μs). For the shortened linker, the dynamics in the open conformation showed similar interconversion rates (relaxation time of 9 ± 7 μs) compared to the timescales in the wt-linker constructs (relaxation time of 15 ± 4 μs).

**Figure 6:**
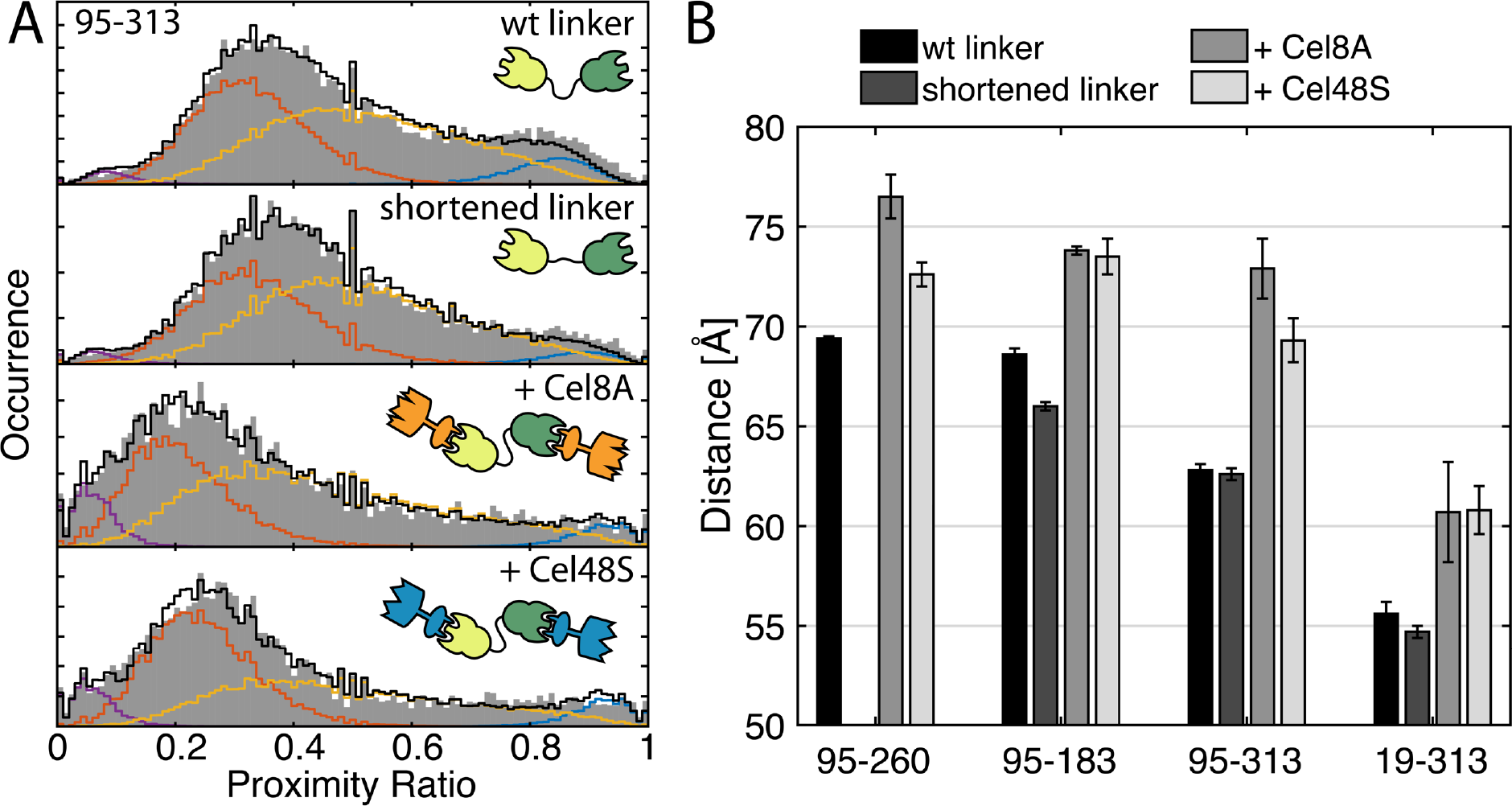
The effect of linker shortening and enzyme binding on the dynamics of CohI_8_-CohI_9_. (A) Dynamic photon distribution analyses (PDA) of the 95-313 construct with wt linker, shortened linker, and wt linker with addition of the cellulosomal enzymes Cel8A and Cel48S. The spFRET histograms of the proximity ratios (uncorrected FRET efficiencies) are plotted as grey bars and the dynamic-PDA fits are shown as black lines. In addition, the low (red) and high (blue) FRET efficiency populations and the population of molecules showing interconversion during the observation time (yellow) are shown. Additional static low-FRET efficiency populations are shown in purple. (B) Comparison of the inter-dye distances of the open conformation (red in A) for the different constructs with wt linker, shortened linker, and wt linker in the presence of the cellulosomal enzymes Cel8A and Cel48S.

In summary, the effect of a shortened linker on the conformational states and dynamics of CohI_8_-CohI_9_ seems to be minor. As expected, the closed conformation was not affected by the linker length. However, shortening of the linker also had no significant effect on the average distance of the open conformation, suggesting that the linker is not fully extended in the wt CohI_8_-CohI_9_, but assumes a more compacted structure due to the observed cohesin-cohesin interactions or via the formation of secondary structure. This is consistent with the results of a SAXS study where the linker length of a chimeric cohesin tandem construct (Scaf4) was systematically varied from 4 to 128 residues (23), revealing that the maximum extension of the construct plateaued already at a linker length of 39 residues. A recent NMR study of the isolated CohI5 module with its 24-residue long linker of similar composition reported a rigid and extended structure of the linker (29). The absence of a second CohI module in that study further indicates that cohesin-cohesin or cohesin-linker interactions may be responsible for the observed compaction of the structure. Interestingly, the stability of the cohesin-cohesin interactions was not affected by the linker length, as the interconversion rates between compacted and extended structures showed no significant change.

### Dockerin-binding shifts the conformational space towards the extended state

The primary *in vivo* function of the Cohl modules is the binding of cellulose-processing enzymes. We investigated the influence of the cellulosomal enzymes Cel8A and Cel48S on the conformational dynamics of CohI_8_-CohI_9_. Both enzymes bind with high affinity to a CohI module through their respective DocI modules with K_D_ values in the range of 10 nM (14). We first confirmed that Cel8A and Cel48S bind to CohI_8_-CohI_9_ under the experimental conditions using FCS (Fig. S12 A). We determined *K_D_* values of 4 nM for Cel8A and 14 nM for Cel48S (Fig. S12 B). In the presence of binding partners, the dynamic behavior of CohI_8_-CohI_9_ persisted (Fig. S13). The dynamic PDA of CohI_8_-CohI_9_ in the presence of Cel8A and Cel48S are shown in Fig. 6A for the 95-313 construct. The results for all four constructs are shown in Fig. S14 and summarized in Fig. 6B and Table S3. For construct 95-313 (Fig. 6A), the average distance in the extended conformation increases from 63 Å in the absence of binding partners to 73 Å and 69 Å in the presence of Cel8A and Cel48S, respectively. This extension of the open state was consistently observed for all constructs (Fig. 6B). Averaged over all constructs, no significant shift of the dynamic equilibrium towards the extended state was observed, with average rates of opening and closing of 1.9 ± 1.5 ms^−1^ and 0.5 ± 0.3 ms^−1^ for Cel8A and 1.7 ± 0.6 ms^−1^ and 0.5 ± 0.1 ms^−1^ for Cel48S, similar to the rates observed in the absence of binding partners of 2.0 ± 0.4 ms^−1^ and 0.5 ± 0.3 ms^−1^.

In summary, the interaction of CohI_8_-CohI_9_ with the cellulosomal enzymes Cel8A and Cel48S resulted in an extension of the open state, while leaving the conformational dynamics largely unaffected. The observed dynamic structure of the CohI_8_-CohI_9_ fragment in complex with Cel8A is supported by SAXS studies on CohI_1_-CohI_2_ with the same enzyme (26) and enzyme-bound hybrid mini-cellulosomes (23) which identified a dynamic and extended structure. Likewise, the fragment CohI_3_-CohI_4_-CohI_5_ in complex with Cel8A has been observed to adopt both extended and compacted structures mediated by cohesin-cohesin interactions (27). The persistence of the compacted conformation in the presence of cohesin-dockerin interactions implies that the dockerin-binding interfaces on the cohesins are not involved in cohesin-cohesin interactions.

### All-atom molecular dynamics simulations provide an atomistic picture of the interactions

From the single-molecule FRET experiments, we could identify a compacted state of the CohI_8_-CohI_9_ fragment. To complement the FRET information provided by the four distances, we performed all-atom molecular dynamics simulations of CohI_8_-CohI_9_ in explicit solvent (see M&M). In total, six trajectories of 200 ns length were simulated from the extended starting configuration, resulting in a different evolution of the system despite the use of identical starting coordinates as evidenced by the root-mean-square deviation (RMSD) of the trajectories (Fig. S15). A detailed display of one of the MD runs is given in Fig. 7A. Snapshots of the trajectory are displayed at the indicated time points above the graph. For the first 50 ns of the simulation, the protein sampled many different “open” structures without forming inter-modular contacts. The two modules then contacted each other and form a stable interaction after ~60 ns, which persisted for the rest of the simulation, showing little additional change in the RMSD (black line in Fig. 7A). As a second parameter for investigating the cohesin-cohesin interactions, we computed the center-of-mass (COM) distance between the two modules (grey line).

**Figure 7:**
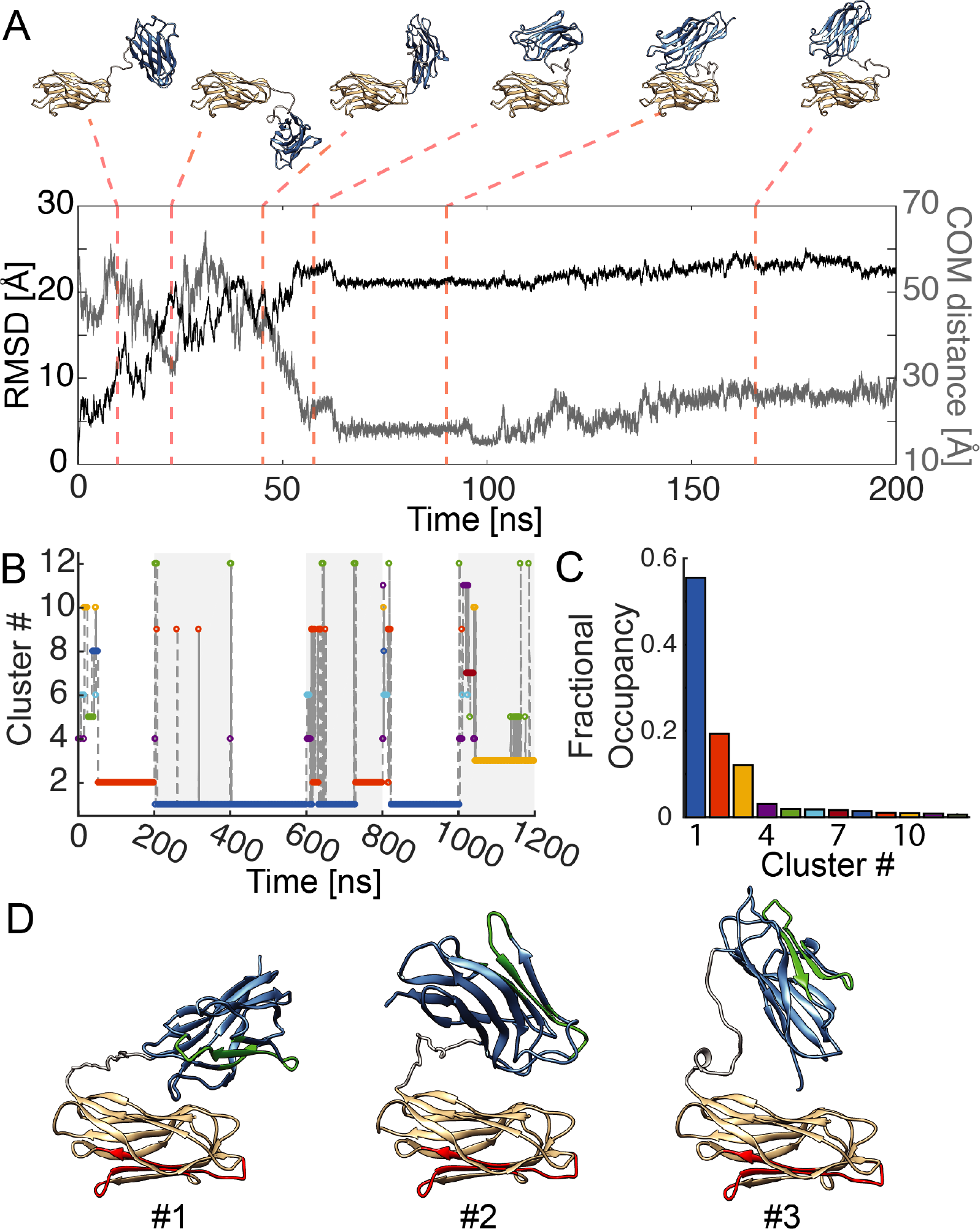
All-atom molecular dynamics simulations of the CohI_8_-CohI_9_ fragment. A) A representative trajectory of a 200 ns MD simulation. Top: Snapshots at the indicated time points of the trajectory. Starting from an extended configuration, the two modules come into close contact after ~60 ns. After the initial contact formation, only minor movements occur without dissociation. Bottom: The root-mean-square deviation (RMSD) with respect to the starting structure plateaus after ~60 ns. Analogously, the center-of-mass (COM) distance between the two modules indicates a close contact. From 100 ns onwards, slight structural rearrangements in the compacted form are evident. B) A cluster analysis of all 6 MD simulations of 200 ns each was performed to obtain global structures of the compacted conformation (shading indicates the individual trajectories). Trajectories 2,3,4 and 5 are dominated by cluster #1. C) Fractional occupancy of the individual clusters. The top three clusters constitute over 80% of the global trajectory. D) The average structures of the clusters 1-3. Regions colored in green and red indicate the binding interfaces for DocI modules (35).

The COM distance contains more specific information about the interaction between the two modules, revealing a slow conformational readjustment from 100 ns till the end of the simulation. During this period, the modules stay in contact, but perform a twisting motion with respect to each other, transitioning from a collinear to a perpendicular orientation. Similar behavior of inter-modular docking during the first half of the simulation was observed for all repeats (Fig. S16). In some cases, detachment of the modules was observed, persisting for only ~10 ns. To obtain an overview of the sampled stable configurations of CohI_8_-CohI_9_, we performed a global cluster analysis over all six trajectories (Fig. 7B-C). The conformational landscape is dominated by three main clusters, accounting for ~85 % of the trajectories (Figure 7D). Clusters 1 and 2 were found in multiple trajectories whereas cluster 3 is unique to trajectory 6. The first cluster adopted a structure showing a perpendicular orientation of the modules. The second and third largest clusters both showed structures where one module contacts the other in a “heads down” orientation, however showing opposite orientation of the top module between cluster 2 and 3. We also performed individual cluster analyses for each trajectory (Fig S17). Each trajectory was dominated by 1 or 2 clusters that accounted for at least 50% of the trajectory. The structures for all major clusters are shown in Fig. S18 for a complete picture of all stable conformations adopted during the simulation. It should be noted that the identified binding modes here are different from the cohesin-cohesin interaction previously observed in crystals (42). To compare the MD simulations with our experimental results on the closed state of CohI_8_-CohI_9_, we determined FRET-average distances from the MD trajectories as described before for the open state (Table S4). Given the high FRET efficiency of the closed state and thus increased uncertainty of the experimentally determined distances, we find reasonable agreement, however the MD-derived distances are consistently larger than the experimental distances by ~8 Å on average. A possible reason could be transient dye-dye interactions that might occur at the short interdye distances, resulting in deviations of the experimentally derived distances (44).

The experimental results of cohesin-cohesin interactions raised the questions whether specific interactions are present, and what role the cohesin-dockerin binding interfaces play. Overall, the simulations show a variety of binding modes. While the global cluster analysis revealed three predominant structures, each of the individual clusters still exhibits some degree of internal variance, indicated by average pairwise RMSD distances within the clusters of 7-8 Å and standard deviations of the RMSD of 2.5-4 Å. This suggests that a multitude of different binding modes are present and the interaction between the modules is not dominated by one specific conformation. To further investigate the cohesin-cohesin interactions, we quantified the number of intermodular contacts formed per residue. Intriguingly, arginine 6 in Cohl_8_ is found to be involved in intermodular contacts in all three major clusters (Fig. S19 A), predominantly forming salt bridges and hydrogen bonds with the residues Q195, T260/E261 or E300/D302 in Cohl_9_ (Fig. S19 B-C). Indeed, in the simulations, the observed conformational space changes significantly when R6 is replaced by a glycine, leading to less frequent formation of stable cohesin-cohesin interactions (Fig. S19 D-E and Fig. S20). While we did not test this mutation experimentally, T260 was mutated to a cysteine and fluorescently labeled in construct 95-260, placing the fluorescent dye in direct vicinity of Q195 and E261. Indeed, in the experiments, this construct showed the lowest population of the closed state (Fig. 4A-D) due to a reduced rate of closing of 0.14 ms^−1^ compared to 0.5-0.9 ms^−1^ for the other constructs (Table 2), indicating that cohesin-cohesin interactions are hindered in this construct. Interestingly, R6 appears in cohesins 4-8, but is absent in cohesins 1-3 and 9, while Q195 is unique to CohI_9_ where it replaces a serine present in the other cohesins (Figure S21). Likewise, E261, E300 and D302 are only found in CohI_9_ and are predominantly replaced by lysine (E261 and E300) or glutamine (D302). The intermodular interactions are thus expected to be different for other cohesin pairs and could potentially serve as a mechanism to fine-tune the quarternary structure of the scaffoldin.

With respect to the enzymatic activity of the cellulosome, it is important to consider the formation of cohesin-cohesin interactions in the context of the accessibility of the dockerin-binding sites. To this end, we colored the residues that have previously been identified to be involved in contacts to DocI modules in Fig. 7D and Fig. S18 in red and green (see Fig. S1 for the colored residues) (35). Interestingly, the DocI-binding surfaces on the CohI modules are mostly unobstructed by the cohesin-cohesin interaction in all stable structures. Hence, the binding of enzymes to the cohesins is not expected to be hindered by cohesin-cohesin interactions. Likewise, the dockerin-cohesin interaction should not interrupt the formation of compacted structures, as is confirmed by our experimental results. Regarding the intermodular linker, we observed no significant formation of stable secondary structure or long-lived interactions within the linker or between the linker and the cohesin domains. In agreement with the experimental data, the linker was rarely extended and often assumed compacted conformations.

In summary, the MD simulations reveal that CohI_8_-CohI_9_ consistently adopts compacted conformations that are stable on the timescale of the simulations (200 ns). A multitude of different binding modes was identified, indicating that the interaction between the modules is not dominated by one specific conformation. In these stable conformations, the dockerin-binding interfaces of the cohesins are exposed to the solvent and thus accessible for enzyme binding. This suggests that cohesin-cohesin interactions and dockerin binding are not mutually exclusive.

## Conclusions

Herein, we characterized the conformational dynamics of CohI_8_-CohI_9_ as a minimal tandem subunit of the scaffoldin protein. Our results show that CohI_8_-CohI_9_ transitions on the millisecond timescale between a flexible extended family of structures and compacted states mediated by cohesin-cohesin interactions. The conformational states and dynamic equilibrium are not influenced by shortening of the inter-cohesin linker. Addition of DocI-containing enzymes preserved the conformational dynamics, but showed a higher cohesin-cohesin distance in the extended state. Molecular dynamics simulations identified potential binding modes for the cohesin-cohesin interaction. All stable conformations identified from the molecular dynamics simulations showed no obstruction of the cohesin-dockerin binding interfaces.

By probing the structural dynamics using four different FRET sensors, we could obtain a detailed understanding of the conformational space of the CohI_8_-CohI_9_ fragment. In principle, the different FRET sensors should yield identical results for the kinetic rates. While we obtained good agreement between the different constructs for most parameters, we also observed some deviations. For example, construct 95-260 showed a significantly reduced transition rate to the closed conformation compared to the other constructs. Based on the MD simulations, we could show that this deviation is likely caused by the close proximity of residue T260 to residues involved in the formation of intermodular contacts. Thus, our study also highlights the importance of testing different labeling position in smFRET experiments to ensure that the fluorescent labeling does not drastically alter the properties of the biomolecule or interfere with its function.

To understand the origin of synergistic effects in cellulosomes, it is essential to obtain a detailed global picture of the structural organization and interactions of the individual functional modules. Our study suggests that cohesin-cohesin interactions might play an essential role for the precise spatial arrangement of the various enzymes. The structure of the cellulosome is inherently dynamic and needs to adapt to changing environments to ensure efficient access of the catalytic units to the crystalline cellulose within the complex mesh of hemicellulose, lignin and pectin. As such, the flexibility of the scaffoldin protein provided by the inter-cohesin linkers is essential for its structural variability. Once the structural rearrangement is completed, however, cohesin-cohesin interactions may be essential for bringing the catalytic subunits into close contact again to provide the high cellulolytic activity through proximity-induced synergistic effects.

Cohesin-cohesin interactions are also an important factor to be considered in the design of artificial mini-cellulosomes for industrial applications. These designer cellulosomes are often chimeras of CohI modules from different organisms to allow control of the enzyme composition through orthogonal cohesin-dockerin interactions. Our results indicate that, in addition to the choice of enzymes, the compatibility of the applied cohesin modules and the flexibility of the linker should be considered to maximize the synergistic effects.

## Acknowledgements

We thank Alvaro H. Crevenna for help with obtaining homology models for the simulations and Sigurd Vogler for performing initial experiments. A.B. and D.C.L. gratefully acknowledge the financial support of the Deutsche Forschungsgemeinschaft through Grants SFB1035 (Projects A11), and support from the Ludwig-Maximilians-Universität through the Center for NanoScience and the BioImaging Network.

## Author contributions

A.B. performed and analyzed single-molecule experiments and molecular dynamics simulations. J.H. performed fluorescent labeling and single-molecule experiments. Y.B. and D.F. expressed and purified proteins. E.B. and D.C.L. designed the experiments and oversaw the project. A.B. wrote the manuscript, with contributions from all authors.

## Materials and Methods

### Protein expression, purification and fluorescent labeling

Protein expression and purification were performed as described previously (45). Proteins contained a His_6_-tag for purification that was not removed. For fluorescent labeling, protein solutions were adjusted to 50 μM and oxygen was removed from the buffer (PBS). Labeling was performed at 10-fold molar excess of the dyes Atto532 and Atto647N (ATTO-TEC GmbH, Siegen, Germany) for 3 hours at room temperature in the presence of 1 mM TCEP. Unreacted dye was removed by ultrafiltration.

### Single-molecule FRET measurements

Single-molecule FRET experiments were performed using a custom-built setup as described previously (34) that combines pulsed interleaved excitation (PIE) (40) with multiparameter fluorescence detection (MFD) (46). With MFD-PIE, it is possible to determine the FRET efficiency, labeling stoichiometry, fluorescence lifetime and anisotropy for both donor and acceptor fluorophores for every molecule. Labeled CohI_8_-CohI_9_ constructs were diluted to a concentration of ~100 pM in buffer containing 25 mM HEPES, 150 mM NaCl and 2 mM CaCl_2_ at pH 7.25. Unlabeled Cel8A or Cel48S was added at concentrations of 50 nM. Excitation powers of 100 μW were used for both donor and acceptor lasers (as measured at the back aperture of the objective). Bursts were identified using a sliding time-window burst search using a time-window of 500 μs and a count rate threshold of 10 kHz(47). Filtering of photobleaching and blinking events was achieved through the ALEX-2CDE filter with an upper limit of 10 (38). Further selection of double-labeled molecules was performed using the stoichiometry parameter with a lower limit of 0.45 and an upper limit of 0.80. Accurate FRET efficiencies were calculated based on the intensities in the donor and FRET channels using correction factors for spectral crosstalk (*α*) of 0.02 and direct excitation of the acceptor fluorophore (*δ*) of 0.06. Differences in the detection efficiencies and quantum yields of the donor and acceptor fluorophores were accounted for using a *γ*-factor of 0.66. The accurate FRET efficiency is then given by (34, 48):

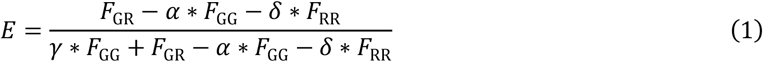

where *F*_GG_, *F*_GR_ and *F*_RR_ are the background-corrected photon counts in the donor channel after donor excitation, the acceptor channel after donor excitation (FRET signal), and the acceptor channel after acceptor excitation, respectively. Burstwise fluorescence lifetimes of the donor and acceptor fluorophore were determined using a maximum likelihood estimator approach (34, 49). For the static FRET lines, the donor lifetime in the absence of the acceptor was determined using donor-only molecules from the measurements selected by a stoichiometry threshold (S > 0.98). Contributions of fast linker fluctuations to the static FRET line were accounted for using a Förster radius, *R*_0_, of 59 Å and an apparent linker flexibility of 5 Å (32). All data analysis was performed using the open-source software package *PAM* written in MATLAB (50).

### Dynamic photon distribution analysis (PDA)

For the photon distribution analysis (31, 51), photon counts from selected single-molecule events were re-binned to equal time bins of 0.25 ms, 0.5 ms and 1 ms length, and histograms of the proximity ratio were computed. The proximity ratio is calculated from the raw photon counts in the donor and FRET channels (*S_D_,S_F_*) by:

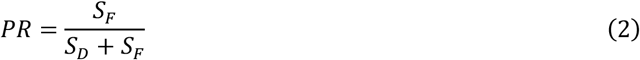

In dynamic PDA, the mixing of states during the fixed observation time as a function of interconversion rates can be solved analytically to describe the dynamic contribution to the observed proximity ratio histogram (32). Due to the dynamic interconversion between different states, the shape of the proximity ratio histogram changes depending on the time bin size. The data were fit using a two-state dynamic model with the addition of one or two minor static low-FRET states. The width of the respective distance distributions, *σ_R_*, was globally fixed at a fraction of the inter-dye distance, *σ_R_* = 0.064 *R*. The proportionality factor was determined from measurements of static double-labeled double-stranded DNA molecules, and thus only accounts for apparent broadening of the distance distribution due to acceptor photophysics (52). This assumption reduces the number of free fit parameters significantly and is justified because no static broadening of the FRET efficiency distribution due to conformational heterogeneity is expected for the studied system. For each dataset, all fit parameters were globally optimized using the proximity ratio histograms obtained for the three different time bin lengths (Figure S5).

### Species-selective fluorescence correlation spectroscopy

Species-selective fluorescence correlation functions were determined as follows: For every burst, photons in a time window of 50 ms around the edges of the burst were added. If another single molecule event was found in the time window, the respective burst was excluded. Correlation functions were calculated for every individual burst and averaged to obtain the species correlation function (53, 54).

For filtered-FCS analysis (33), microtime patterns for the low and high FRET efficiency species were obtained from subpopulations of the experiments directly using FRET efficiency thresholds (Fig. S7A). Donor and FRET-induced acceptor decays were stacked, and filters were generated separately for the parallel and perpendicular detection channels (Fig. S7B-D), which were cross-correlated to circumvent the dead time of the TCSPC hardware and detectors. In this way, the fFCS correlation functions can be calculated down to a limit of 40 ns given our hardware configuration.

FRET-FCS and filtered-FCS curves were fit to a standard single-component diffusion model with up to two kinetic terms, given by:

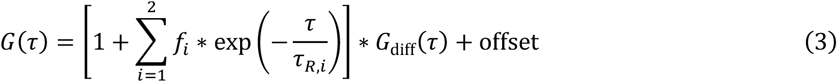

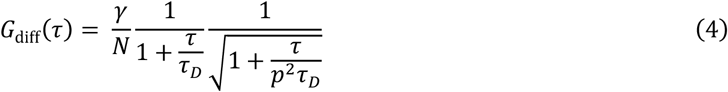

*G*_diff_(*τ*) is the diffusion part of the correlation function, where *N* is the average particle number in the confocal volume, 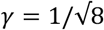 is a geometric factor that accounts for the Gaussian shape of the observation volume, *τ_D_* is the diffusion time and *p* is the ratio of the axial and lateral size of the confocal volume. The amplitudes of the kinetic terms are given by *f_i_* and the relaxation times by *τ_R,i_*. For FRET-FCS, the three correlation functions (donor x donor, FRET x FRET and donor x FRET) were fit using a single kinetic term by globally linking the parameters of the diffusion term and the relaxation time of the kinetic term *τ_R_*, and letting the amplitude of the kinetic term assume negative values for the cross-correlation function. For filtered-FCS, the four correlation functions between the two species A and B (AxA, AxB, BxA, BxB) were fit using two kinetic terms by globally linking the parameters of the diffusion term (with exception of the particle number *N*) and the relaxation times of the kinetic terms *τ_R,i_*, and letting the amplitudes of the kinetic terms *f_i_* for the cross-correlation functions assume negative values.

### Molecular dynamics simulations

A homology model for the two CohI modules was built using SWISS-MODEL (55–58) based on the crystal structures of CohI_9_ (PDB: 3KCP) (42) for the CohI_9_ module, and the crystal structure of CohI_7_ (PDB: 1AOH) (59) for the CohI_8_ module, with sequence similarities of 98.68% and 95.83%, respectively. Torsion-angle rigid-body MD simulations were performed using the Crystallography and NMR system (CNS) (60–63). After addition of the linker peptide and relaxation of the structure, an 80 ns trajectory of the dimer was simulated at 300 K with a time step of 5 fs, treating the cohesin modules as rigid bodies while leaving the covalent bonds in the linker free to rotate. Since the aim of the simulation was to determine equilibrium distances of the extended state, no explicit solvent is included. Every 10 ps, possible positions of the fluorophores were determined using accessible volume (AV) calculations with standard parameters for Atto647N using the FPS software package (36). From the accessible volumes, expected average FRET efficiencies, 〈*E*〉(*t*), are calculated at every time step, *t*, by averaging over all possible combinations of donor and acceptor positions using a Förster *R*_0_ radius of 59 Å. The FRET efficiencies, 〈*E*〉(*t*), are averaged over all time steps and converted back to distances, yielding the FRET-averaged expected distances of the simulation. Error bars are determined by bootstrapping.

All-atom MD simulations were performed with the AMBER16 molecular dynamics package using the ff14SB force field (64). The molecule was solvated in a pre-equilibrated box of TIP3P water using a truncated octahedron geometry with a minimum distance between solute and the periodic boundaries of 2 nm. The charge of the system was neutralized by addition of 23 sodium ions. Two additional sodium and chloride ions were added, resulting in an excess salt concentration of 2 mM. Initial energy minimization of the extended starting structure was performed using the steepest descent method for 10 steps followed by 190 steps using the conjugate gradient method. For equilibration, the system was heated to 298 K over 50 000 steps with a step size of 2 fs at constant volume, and subsequently run for 50 000 additional steps at 298 K with pressure scaling enabled. Individual MD runs were performed for at least 100 ns at 2 fs step size using the NPT ensemble with the Monte Carlo barostat. Trajectories were written at a resolution of 10 ps. The individual trajectories were obtained using the same equilibrated starting structure with random assignment of the initial velocities. On a single Nvidia GTX 1080 Ti GPU, the simulation typically ran at 50 ns a day. Analysis of the MD trajectories was performed using the *cpptraj* utility of the AMBER16 software package (65). Clustering was performed using the hierarchical agglomerative algorithm using the average-linkage criterion and a cluster number of 12. Structural figures were generated using UCSF Chimera (66).

